# Germination-stage transcriptomics of the grapevine rust fungus *Neophysopella tropicalis* reveals protein effectors with plant immunity-suppressing activity

**DOI:** 10.1101/2025.10.22.683938

**Authors:** Thiago Maia, Joao V. F. Mazzei, Renato G.H. Bombardelli, Claudia B. Monteiro-Vitorello

**Author notes:** Corresponding authors: (TM); (CBM-V).

## Abstract

Germination of urediniospores represents a critical early stage in the life cycle of *Neophysopella tropicalis*, the causal agent of Asian grapevine leaf rust in Brazil. Rust fungi deploy diverse effector proteins to manipulate host immunity and establish infection. This study identified and functionally characterized secreted proteins expressed during early fungal development using RNA-seq analysis of in-vitro-germinated urediniospores. Effectorome analysis revealed 386 putative secreted proteins with hallmarks of fungal effectors, designated as *N. tropicalis* effector candidates (NtECs). Fifteen highly expressed *NtECs* were selected for expression profiling by RT-qPCR across infection stages. Fluorescence microscopy with WGA-FITC staining defined key phases of pathogenesis. Although identified during germination, seven *NtECs* showed elevated transcript accumulation during the penetration and early biotrophic phases, suggesting roles in host entry and establishment. To evaluate immune-suppressing activity, nine NtECs were delivered into *Nicotiana benthamiana* using the Type III Secretion System of *Pseudomonas fluorescens* EtHAn. Three effectors (NtEC-05, -09, and -10) strongly suppressed AvrB-triggered cell death, and three others (NtEC-11, -12, and -13) showed moderate suppression. This work provides the first characterization of *N. tropicalis* effectors, offering insights into rust pathogenesis and supporting future effector-informed strategies for grapevine resistance.

**Highlight:** The grapevine rust fungus *Neophysopella tropicalis* deploys effectors whose expression aligns with early infection and immune suppression, revealing key virulence mechanisms and guiding effector-informed strategies for grapevine rust resistance.

## Introduction

Grapevine (*Vitis* spp.) ranks among the world’s most important fruit crops and, in recent years, has gained increasing prominence in Brazil (de Oliveira et al., 2020). For fresh fruit consumption, the American *Vitis labrusca* cultivars predominate in Brazilian vineyards, mainly due to their vegetative traits well adapted to the country’s environmental conditions (Santos et al., 2020a; da Silva et al., 2022). However, maintaining this crop requires significant investment in integrated disease management combining cultural and chemical control (Gramaje et al., 2018). The problem is especially acute for *V. labrusca*, which is vulnerable to a wide range of filamentous plant pathogens (Primiano et al., 2017; Santos et al., 2020b). Among these, *Neophysopella tropicalis*, the causal agent of Asian grapevine leaf rust (AGLR), stands out as one of the most destructive threats to the crop (Rasera et al., 2023). AGLR was initially attributed to species within the genus *Phakopsora*, but has since been reassigned to the genus *Neophysopella* based on morphological and phylogenetic analyses (Ji et al., 2019).

A survey of Brazilian *Neophysopella* isolates associated with AGLR revealed that all 50 monouredinial cultures from the south-central region were identified as *N. tropicalis*, whereas six isolates from the northeast were classified as *Neophysopella meliosmae-myrianthae* (Santos et al., 2020a). Therefore, *N. tropicalis* is the predominant causal agent of AGLR in Brazil and the sole species found in the southern and southeastern regions, which together account for 75% of the nation’s grape production (IBGE, 2024). Despite its high prevalence and destructive potential, the molecular basis of *N. tropicalis* virulence remains unexplored, particularly the effector proteins deployed during early infection stages that may suppress host immunity and facilitate colonization.

A hallmark of rust fungi is their large secretome, encompassing hundreds of predicted genes encoding secreted proteins, including effectors relevant to establishing infection and manipulation of the host immunity and physiology (Lorrain et al., 2019; Figueroa et al., 2021). In *N. tropicalis*, urediniospore germination represents the initial and critical step in pathogenesis, marking the onset of infection. Understanding the molecular events occurring during this stage is thus essential to elucidate how filamentous pathogens penetrate and colonize their hosts (Shen et al., 2017; He et al., 2020).

To investigate these early infection processes, germinating structures from obligate biotrophic pathogens grown in vitro provide valuable material for RNA sequencing (RNA-seq) studies, particularly when public reference genomes are unavailable. This approach allows isolation of pathogen RNA free from plant biomass. Moreover, candidate effector genes identified from cDNA libraries constructed from germinating urediniospores frequently exhibit transcript accumulation in planta between 24 and 72 hours post-inoculation (Maia et al., 2017), confirming that germlings serve as a valuable biological source for identifying effectors induced during plant infection.

In this study, we developed a robust RNA-seq strategy using germinating urediniospores of *N. tropicalis* to identify candidate effector genes expressed at early infection stages and functionally characterize their role in suppressing plant immunity. Our objectives include (i) pinpointing genes encoding secreted proteins during urediniospore germination, (ii) profiling their expression across infection stages, (iii) associating effector expression with specific pathogenesis phases, and (iv) establishing a protocol for transient expression of *N*. *tropicalis* secreted proteins in planta leaves using the Type III Secretion System of *Pseudomonas fluorescens* EtHAn. This comprehensive investigation fills a critical knowledge gap in *N. tropicalis* biology, providing novel insights into the molecular mechanisms underlying its pathogenicity and supporting future efforts to develop grapevine cultivars resistant to this destructive disease. Notably, no candidate effector sequences from *N. tropicalis* are currently available in GenBank or other public databases, highlighting the urgent need for genomic and transcriptomic studies to uncover its virulence factors.

## Material and methods

### Urediniospores germination

Urediniospores from the single-pustule isolate Aglr64 of *N*. *tropicallis* (Jales-SP), kindly provided by the group of Professor Lilian Amorim (ESALQ-USP) and previously molecularly characterized by Santos et al. (2020a), were inoculated onto the abaxial surface of fully expanded, healthy leaves of *Vitis labrusca* cv. Niagara Rosada to promote multiplication under controlled growth chamber conditions, following Rasera et al. (2023). Ten days after inoculation, leaves exhibiting a high density of sporulating pustules were harvested, and approximately two grams of fresh urediniospores were collected by gently brushing the abaxial surface of infected leaves. The urediniospores were then washed with sterile distilled water containing 0.05% Tween 80 (Sigma-Aldrich), evenly distributed onto polystyrene Petri dishes with a thin film of sterile water, and incubated in darkness at 22°C. Germination was confirmed by light microscopy after 16 hours of incubation. The resulting *N. tropicalis* germlings were collected, flash-frozen in liquid nitrogen, and stored at –80°C until total RNA extraction.

### RNA preparation and RNA-seq library for sequencing

Total RNA was extracted from germinated urediniospores using RNeasy Plant Mini Kit (Qiagen), according to the manufacturer’s instructions. Any contaminating of genomic DNA was eliminated from the extracted RNA using RNAse-free DNAse I, following the manufacturer’s protocol (Sigma-Aldrich). RNA quality was assessed by a 2100 Bioanalyzer RNA Nano Chip (Agilent). Library construction was generated using the Illumina TruSeq^TM^ Stranded mRNA Library Preparation Kit TruSeq (Illumina) and then paired-end sequenced on a NextSeq 500 platform (Illumina) at Functional Genomics Center, ESALQ, University of São Paulo (Piracicaba, São Paulo, Brazil).

### *De novo* transcriptome assembly and candidate effector proteins prediction

FastQC v0.11.5 was applied to check the quality of the reads (Andrew, 2010). Then, we conducted the trimming and quality filtering using Cutadapt v1.18 (Martin, 2011) to remove the adapters and keep only the reads larger than 50 bp, with no “N” bases, and Phred quality score higher than 30 on average. For the assembly of the transcriptome, we used Trinity v2.14 (Grabherr et al., 2011) with default parameters. The input used was the filtered FastQ files from the sequencing, biological triplicate. As this study focused on the identification of candidate effectors, we ran SignalP v4.1 (Petersen et al., 2011), TMHMM v2.0 (Krogh et al., 2001), and EffectorP v2.0 (Sperschneider et al., 2018) to predict the candidate effectors from *N*. *tropicalis*. We mapped the reads against the *de novo* transcriptome assembly using Hisat2 v.2.2.1 software (Kim et al., 2015) and generated a count table using the FeatureCounts from subread package v2.0.6 (Liao et al., 2014) to estimate the most expressed candidate effector genes in the germinated urediniospores from *N*. *tropicalis*. Subsequently, the final set of predicted effectors was annotated using BlastP against the nr-NCBI protein database.

### Selection and validation of the candidate effector genes using RT-PCR

We selected the fifteen most highly expressed candidate effector genes with complete open reading frames (ORFs) to validate the bioinformatic predictions through reverse transcription polymerase chain reaction (RT-PCR). For cDNA synthesis, 1.0 µg of total RNA extracted from germinated urediniospores of the pathogen was used as input, with the Oligo(dT)12–18 primer and the SuperScript First-Strand Synthesis System (Thermo Scientific) for RT-PCR kit. Gene-specific primers were designed to amplify the ORF sequences, excluding the signal peptide, using the synthesized cDNA as a template. PCR amplification was performed using the Phusion High-Fidelity PCR Master Mix (Thermo Scientific). The resulting amplicons were subsequently used for cloning and downstream functional analyses (see section 4.7).

### Expression analysis of *N*. *tropicalis* candidate effector genes using RT-qPCR

For RT-qPCR assays, inoculations were performed using a suspension of 1 × 10 urediniospores mL ¹, following Rasera et al. (2023). Total RNA was extracted from rust-infected grapevine leaves of *V. labrusca* cv. Niagara Rosada collected at 24, 48, and 72 h post inoculation (hpi) and at 7 days post inoculation (dpi), following a CTAB-based protocol (Iandolino et al., 2004). For each time point, three biological replicates, each consisting of independently infected leaves, were collected. Additionally, RNA from dormant and germinated urediniospores was isolated using the RNeasy Plant Mini Kit (Qiagen). Following DNase treatment, 1 µg of RNA from each sample was used for cDNA synthesis with the SuperScript First-Strand Synthesis System for RT-PCR (Invitrogen), using the Oligo(dT)12–18 primer in a final reaction volume of 20 µl. The quality and efficiency of cDNA synthesis were evaluated using endogenous genes from both the pathogen and the host (Supplementary Fig. S4).

Relative gene expression was calculated using the 2^-ΔΔCt^ method (Livak and Schmittgen, 2001), based on the average Ct values from three biological replicates and two technical replicates per time point. RT-qPCR reactions were carried out using the GoTaq RT-qPCR Kit (Promega) on a 7300 PCR System (Applied Biosystems), with the following cycling conditions: initial denaturation at 95 °C for 2 minutes, followed by 40 cycles of 95 °C for 15 seconds and 60 °C for 60 seconds. Amplification specificity was confirmed by melting curve analysis, performed from 60 to 95 °C with a temperature increment of 1 °C every 30 seconds.

### Microscopy analyses of *N*. *tropicalis* infection stages

Fluorescence microscopy was performed using a Leica DM4000 microscope. Fungal hyphae were visualized by staining with 20 μg/mL wheat germ agglutinin conjugated to fluorescein isothiocyanate (WGA-FITC; Sigma-Aldrich), following the protocol described by (Marrafon-Silva et al., 2024). FITC fluorescence was excited at 488 nm, and emission was detected within 500–550 nm range. To observe *N. tropicalis* infection structures, rust-inoculated grapevine leaves were collected at 24, 48, and 72 hpi, as well as at 5, 7, and 9 dpi.

### Cloning the *N*. *tropicalis* effector candidate genes into pEDV6 vector

*NtEC* sequences were cloned into the pEDV6 vector (Maia et al., 2017) using the Gateway^®^ LR Clonase™ II Enzyme Mix kit (Invitrogen), following the manufacturer’s instructions. The in-frame fusion and sequence integrity of all pEDV6::*NtEC* constructs were verified by sequencing. *Escherichia coli* DH5α was used for propagation of all pEDV6::*NtEC* constructs. The pEDV6 plasmids expressing *NtECs* were transferred from *E. coli* DH5α to *P. fluorescens* EtHAn by standard triparental mating, using *E. coli* HB101 (pRK2013) as the helper strain (Maia et al., 2022). Empty pEDV6 vectors were maintained in *E. coli* DB3.1. Transconjugated *P. fluorescens* cells carrying the pEDV6::*NtEC* constructs were selected on solid Luria–Bertani (LB) medium supplemented with chloramphenicol (30 μg mL ¹) and gentamicin (25 μg mL ¹).

### Cloning the *N*. *tropicalis* effector candidate genes into pEDV6 vector

To assess whether the NtECs can suppress plant immunity response, we employed an AvrB-induced cell death suppression assay (Maia et al., 2022). Suspensions of *P. fluorescens* EtHAn carrying pVSP6::*AvrB* (Innes et al., 1993) and *Pf* EtHAn carrying pEDV6::*NtECs* were mixed in a 1:1 ratio to a final concentration of 2 × 10 CFU ml ¹ and co-infiltrated into leaves of 4–5-week-old *N. benthamiana* plants using a needleless syringe. Co-infiltrations of *Pf* EtHAn (pVSP6::*AvrB*) with *Pf* EtHAn (pEDV6 empty vector) served as negative controls. Cell death suppression was considered to occur when an effector candidate reduced or prevented tissue collapse in at least 40% of the co-infiltrated leaves.

### Structural prediction of *N. tropicalis* effectors

Three-dimensional structures of NtECs with plant immune suppression activity (excluding signal peptides) were predicted using AlphaFold v3 (Abramson et al., 2024) using default settings, selecting the top-ranked of five models (model_0.cif). High-confidence structures (pTM > 0.5) were compared to known proteins via FoldSeek (van Kempen et al., 2024), aligned using US-align (Zhang et al., 2022), and visualized in PyMOL v3.1 (Schrödinger and DeLano, 2020).

## Results

### Urediniospore germination and *de novo* transcriptome assembly

Fresh urediniospores of *N*. *tropicalis* harvested from *Vitis labrusca* leaves (Fig. 1A), were washed and placed to germinate on polystyrene plates. After 16 hours of incubation, 88% of the urediniospores had developed long germ tubes (Fig. 1B, C). Germinated urediniospores were then collected for total RNA extraction (Supplementary Fig. S1) and subsequent RNA-seq library construction.

**Fig. 1:**
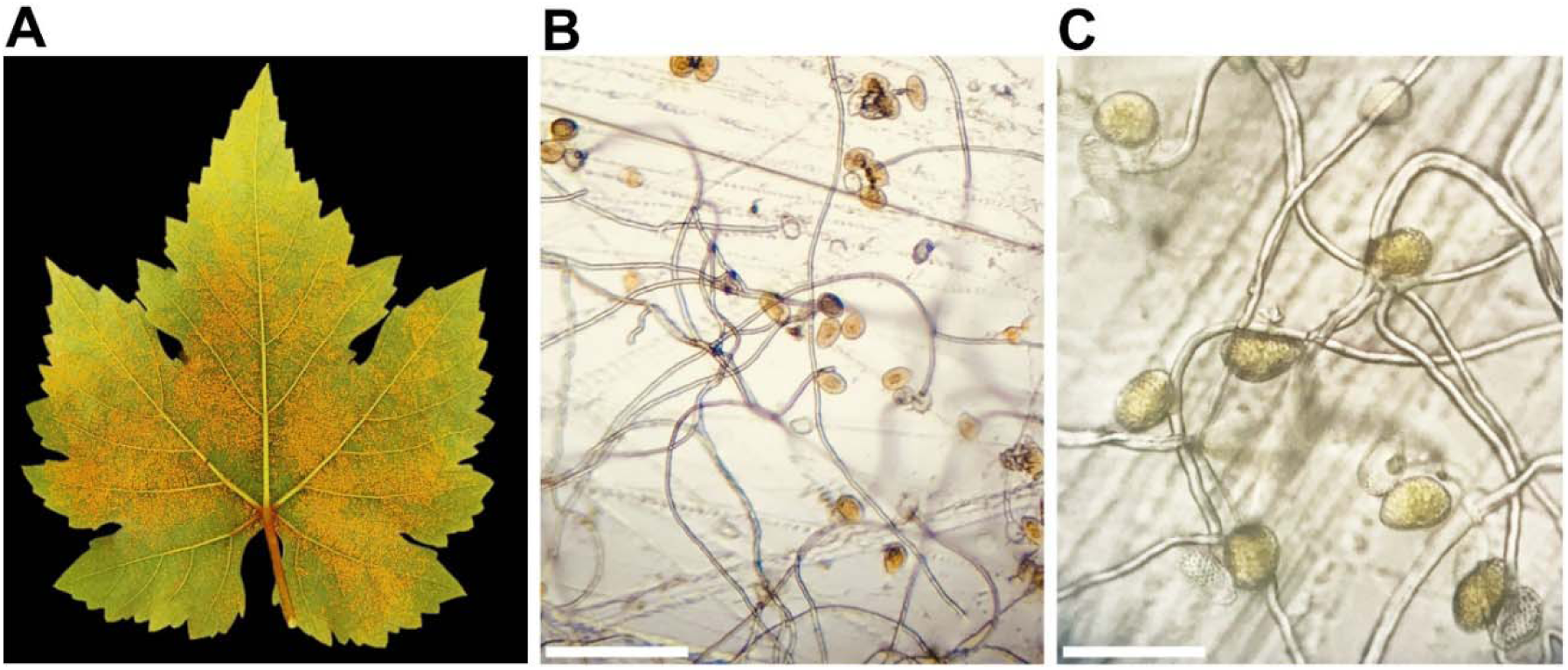
Asian grapevine rust fungus *Neophysopella tropicalis*. (A) Uredinial pustules of *N*. *tropicalis* isolate Aglr64 developing on the abaxial surface of *Vitis labrusca* cv. Niagara Rosada leaves. (B) Light microscopy image of germinated urediniospores and elongating germ tubes on polystyrene after 16 hours of incubation at 22 °C. Scale bar = 120 µm. (C) Close-up view of germ tube emergence from urediniospores on the plate surface. Scale bar = 30 µm.

High-throughput sequencing yielded over 90 million trimmed reads, which were assembled into contigs using the Trinity assembler. The final assembly consisted of 64,333 transcripts, of which 28,968 (45.03%) exceeded 1,000 base pairs in length (Supplementary Table S1). BUSCO analysis revealed that 89.6% of the transcripts showed similarities with fungi sequences deposited in the fungi_odb10 database.

Among the assembled sequences, transcripts DN4595_c0_g1_i2 and DN303_c3_g2_i3 encoded full-length open reading frames (ORFs) with strong similarity to β-tubulin. DN4595_c0_g1_i2 (*Nt*β*Tub1*) shared 100% identity with a known *N. tropicalis* β-tubulin gene (GenBank accession UQJ88286.1), whereas DN303_c3_g2_i3 (*Nt*β*Tub2*) represents a putative novel β-tubulin gene not previously reported in this species, suggesting the existence of a second β-tubulin isoform.

To assess their conservation, we aligned the deduced amino acid sequences of *Nt*β*Tub1* and *Nt*β*Tub2* to their respective top BLASTP hits (Supplementary Fig. S2). In addition, two other contigs, DN600_c1_g1_i2 and DN600_c1_g1_i4, encoded full-length ORFs with high similarity to glyceraldehyde-3-phosphate dehydrogenase (*GAPDH*). These two sequences likely correspond to allelic variants of a single-copy gene, as they differ by only one non-synonymous substitution (Supplementary Fig. S3), consistent with reports of single-copy *GAPDH* genes in other filamentous fungi (Lima et al., 2009). Notably, no *GAPDH* sequence from *N. tropicalis* is currently available in the NCBI GenBank database. These full-length β*-tubulin* and *GAPDH* sequences identified here will serve as endogenous reference genes for *N. tropicalis* to normalize the relative expression of candidate effector genes in forthcoming experiments.

### Candidate effectors identification from germinated urediniospores

From those 64,333 sequences identified in the transcriptome of germinated urediniospores from *N*. *tropicalis*, 6081 (9.45%) encoding for predicted secreted proteins without transmembrane domains. Of those, 386 sequences were classified as candidate effectors using the EffectorP-2.0 (*P* > 0.8). We selected the fifteen most expressed *N*. *tropicalis* effector candidate genes (*NtECs*) based on the higher number of reads mapped per transcript identified in the RNA-seq library (Table 1).

**Table 1:** Fifteen *Neophysopella tropicalis* effector candidate genes (*NtECs*) selected from the germinated urediniospores transcriptome library based on their high transcript abundance (read count per contig).

| *NtEC* | Contig ID | ORF length | Mature ORF | No. of Reads | Protein Size | SignalP 6.0* |
| --- | --- | --- | --- | --- | --- | --- |
| *NtEC*-01 | DN7996_c0_g1_i12 | 480 bp | 426 bp | 22,927 | 160 aas | 0.981 |
| *NtEC*-02 | DN490_c0_g1_i5 | 498 bp | 444 bp | 14,308 | 166 aas | 0.9996 |
| *NtEC*-03 | DN4194_c0_g3_i1 | 471 bp | 402 bp | 11,488 | 157 aas | 0.9977 |
| *NtEC*-04 | DN62_c0_g1_i24 | 681 bp | 624 bp | 8,101 | 227 aas | 0.9695 |
| *NtEC*-05 | DN470_c0_g1_i12 | 384 bp | 324 bp | 3,505 | 128 aas | 0.9997 |
| *NtEC*-06 | DN62_c0_g1_i8 | 540 bp | 480 bp | 2,305 | 180 aas | 0.8636 |
| *NtEC*-07 | DN2816_c0_g2_i23 | 531 bp | 420 bp | 1,287 | 177 aas | 0.9941 |
| *NtEC*-08 | DN1789_c0_g1_i7 | 693 bp | 639 bp | 1,098 | 231 aas | 0.9997 |
| *NtEC*-09 | DN8063_c0_g1_i13 | 474 bp | 408 bp | 1,066 | 158 aas | 0.9998 |
| *NtEC*-10 | DN507_c0_g1_i2 | 378 bp | 318 bp | 996 | 126 aas | 0.9993 |
| *NtEC*-11 | DN2062_c0_g1_i5 | 486 bp | 414 bp | 968 | 162 aas | 0.9998 |
| *NtEC*-12 | DN7996_c0_g1_i17 | 480 pb | 429 bp | 869 | 160 aas | 0.9998 |
| *NtEC*-13 | DN3898_c0_g1_i2 | 306 pb | 246 bp | 761 | 102 aas | 0.9998 |
| *NtEC*-14 | DN1304_c0_g1_i6 | 360 pb | 294 bp | 743 | 120 aas | 0.9998 |
| *NtEC*-15 | DN7682_c0_g3_i1 | 519 bp | 459 bp | 715 | 173 aas | 0.9995 |

\*Signal peptide predicted using SignalP 6.0 (Teufel et al., 2022).

Of the fifteen *NtECs* selected for further characterization, seven (*NtEC-01*, *NtEC-07*, *NtEC-09*, *NtEC-11*, *NtEC-13*, *NtEC-14*, and *NtEC-15*) showed no similarity to any sequences currently available in the NCBI non-redundant (nr) protein database, suggesting that they may represent species-specific genes unique to *N*. *tropicalis* (Table 2). Four candidates (*NtEC-03*, *NtEC-05*, *NtEC-08*, and *NtEC-12*) were homologous exclusively with sequences from rust fungi, indicating a potential specificity to the order Pucciniales. Three candidates (*NtEC-02*, *NtEC-04*, and *NtEC-06*) showed taxonomically restricted similarity to members of the Basidiomycota phylum. Notably, *NtEC-10* exhibited broader cross-phyla conservation, with significant similarity to sequences from both Basidiomycota and Ascomycota, pointing to a more evolutionarily conserved effector candidate (Table 2).

**Table 2:**
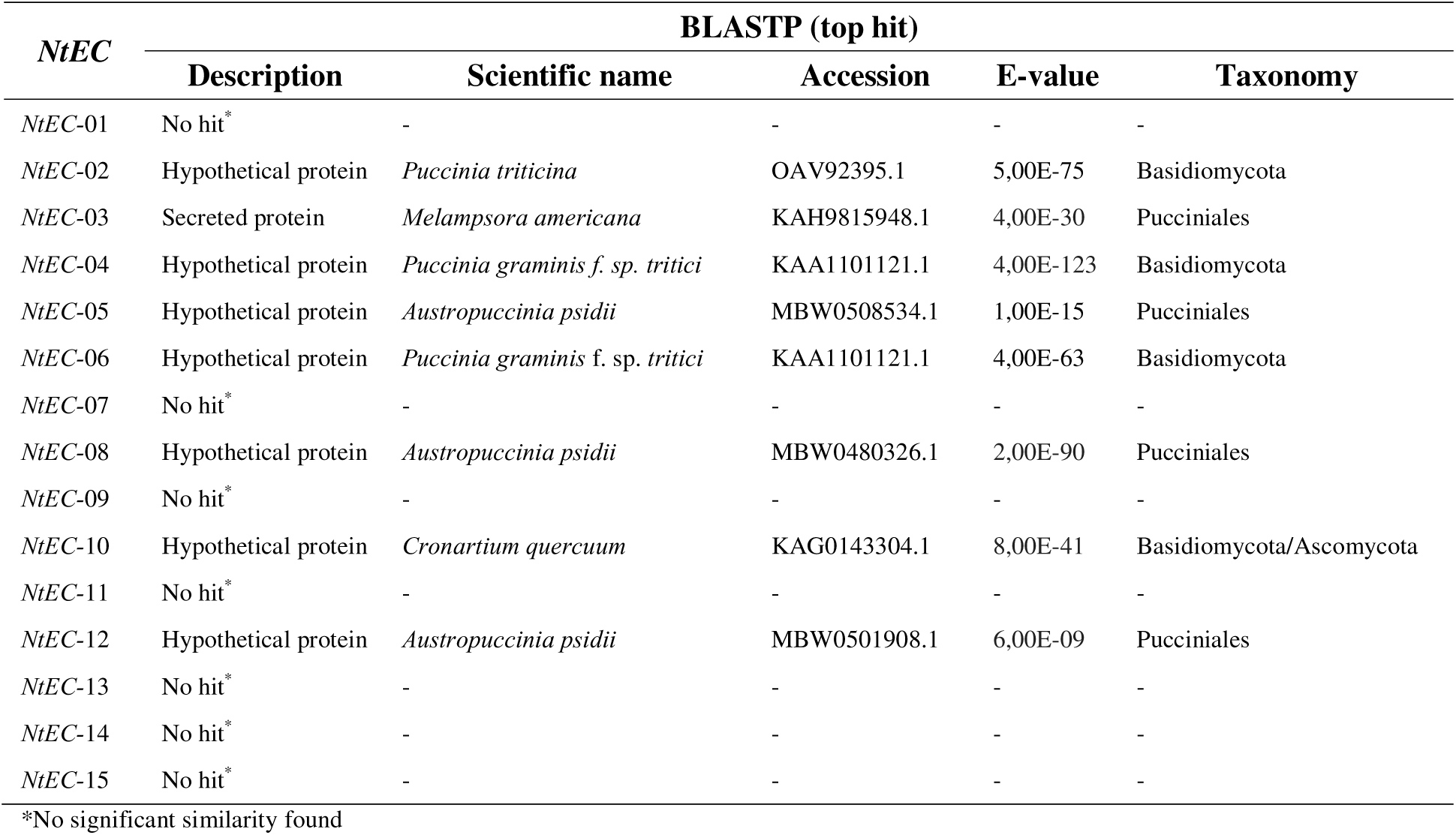
BLAST-based annotation of fifteen *Neophysopella tropicalis* effector candidate genes (*NtECs*) using the NCBI non-redundant (nr) database.

To experimentally validate the bioinformatic predictions of the fifteen *N. tropicalis* effector candidate sequences, we conducted reverse transcription polymerase chain reaction (RT-PCR) assays using complementary DNA (cDNA) synthesized from total RNA extracted from germinated urediniospores. Primers were designed to amplify the coding region of the mature protein, excluding the N-terminal signal peptide sequence predicted by SignalP 6.0. Fourteen of the fifteen *NtECs* produced amplicons of the expected size, consistent with the *in silico* predictions of mature ORFs (Fig. 2; Table 1). One exception was *NtEC-06*, which generated an amplicon of approximately 380 bp instead of the predicted 480 bp. This corresponds to an overall validation accuracy of 93.33% for our computational predictions. The fourteen successfully validated *NtECs* were further subjected to gene expression profiling analyses during the pre-biotrophic and biotrophic developmental stages of *N. tropicalis* to investigate their potential involvement in distinct phases of pathogenesis.

**Fig. 2.**
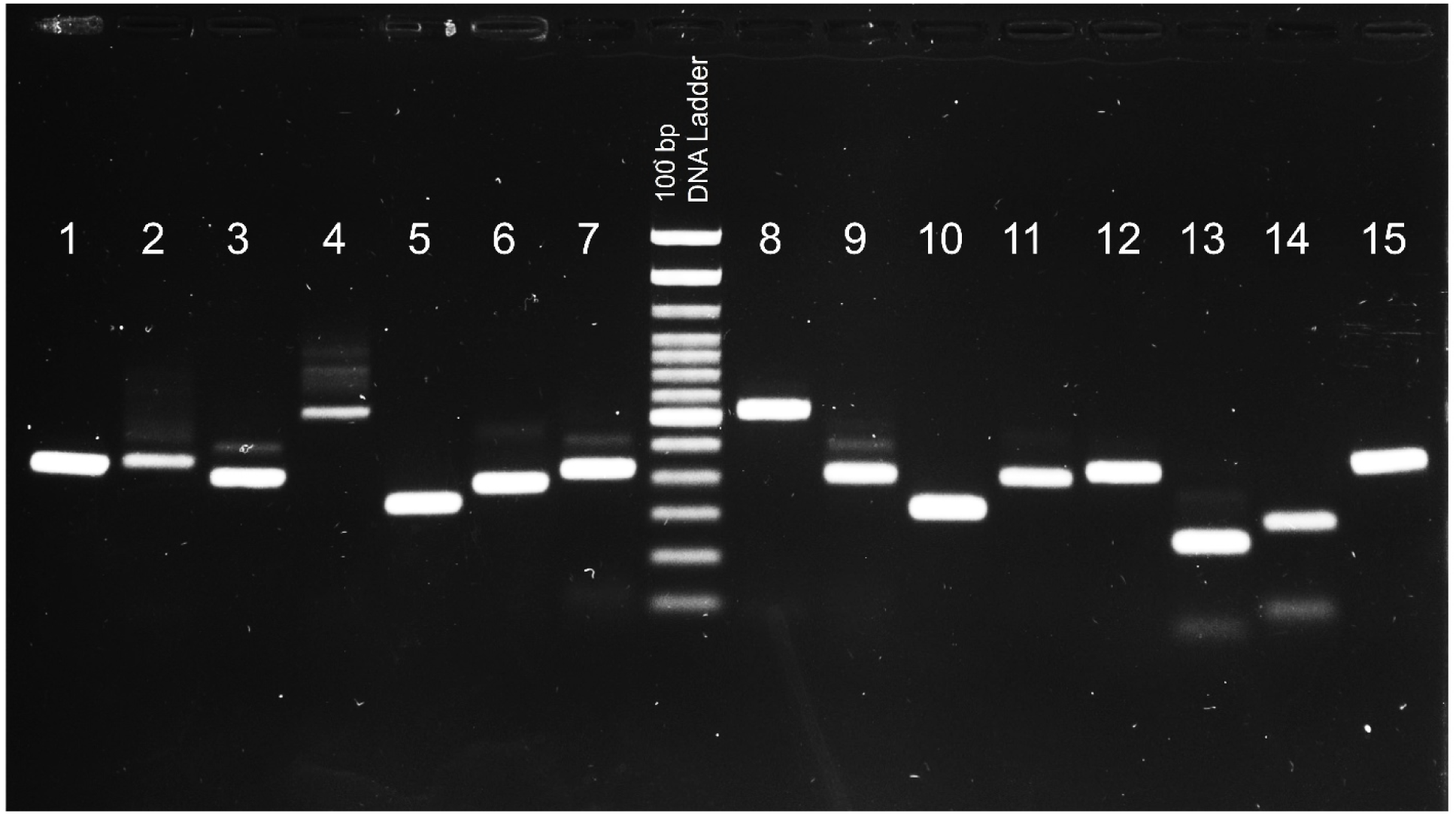
Amplification of fifteen *Neophysopella tropicalis* effector candidate genes (*NtECs*) from germinated urediniospores by reverse transcription polymerase chain reaction (RT-PCR). Agarose gel electrophoresis (1.2%) showing RT-PCR products obtained using cDNA synthesized from RNA extracted from germinated urediniospores as template. Each amplicon corresponds to the mature open reading frame of the respective effector candidate gene, excluding the N-terminal signal peptide sequence. Lane assignments: 1: *NtEC-01*, 2: *NtEC-02*, 3: *NtEC-03*, 4: *NtEC-04*, 5: *NtEC-05*, 6: *NtEC-06*, 7: *NtEC-07*, 8: *NtEC-08*, 9: *NtEC-09*, 10: *NtEC-10*, 11: *NtEC-11*, 12: *NtEC-12*, 13: *NtEC-13*, 14: *NtEC-14*, 15: *NtEC-15*. The primer sequences are listed in Table S2.

### Expression profiles of candidate *Neophysopella tropicalis* effectors

We analyzed transcript accumulation of *N. tropicalis* effector candidate genes (*NtECs*) at distinct stages of the grapevine rust fungus life cycle. To this end, cDNA was synthesized from RNA extracted from both dormant and germinated urediniospores, as well as from infected grapevine leaves collected at 24, 48, and 72 hours post-inoculation (hpi), and 7 days post-inoculation (dpi). To verify the quality of the cDNA derived from these samples, RT-PCR was performed using primers targeting conserved fungal endogenous genes—*Nt*β*Tub1* and *Nt*β*Tub2* (β-tubulin), *NtGAPDH* (glyceraldehyde-3-phosphate dehydrogenase), and *NtCytIII* (cytochrome c oxidase subunit III), alongside the grapevine reference gene *VvActin7* (Supplementary Fig. S4). All fourteen *NtECs* yielded a single amplification product in samples from urediniospores and/or germinated urediniospores. Notably, some *NtECs* were also amplified in infected leaf tissues, suggesting that a subset of these genes may be expressed during the biotrophic phase of infection (Supplementary Fig. S4).

To further characterize their expression, we performed real-time quantitative PCR (RT-qPCR). The *NtECs* exhibited distinct expression patterns throughout the infection process and were classified into six groups based on their peak transcript accumulation (Fig. 3). Among them, *NtEC-08* showed transcript accumulation in both dormant and germinated urediniospores (Fig. 3A). *NtEC-07* displayed a pronounced expression peak in germinated urediniospores and at 24 hpi, suggesting a role during the initial penetration phase (Fig. 3B). Six effector candidates (*NtEC-02*, *-03*, *-10*, *-04*, *-11*, and *-13*) were highly expressed exclusively in germinated urediniospores, indicating potential involvement in pre-biotrophic development (Fig. 3C). *NtEC-01* and *NtEC-14* exhibited peak transcript accumulation restricted to 24 hpi, supporting their likely role in host penetration (Fig. 3D). Notably, *NtEC-05* and *NtEC-12* reached maximum expression at both 24 and 48 hpi, suggesting broader activity across early infection stages (Fig. 3E). Finally, *NtEC-09* and *NtEC-15* displayed peak transcript levels at 48 hpi, possibly reflecting a role in sustaining biotrophic interaction (Fig. 3F). To corroborate the stage-specific expression of the effector candidate genes during *N*. *tropicalis* pathogenesis, we performed a microscopic time-course analysis.

**Fig. 3.**
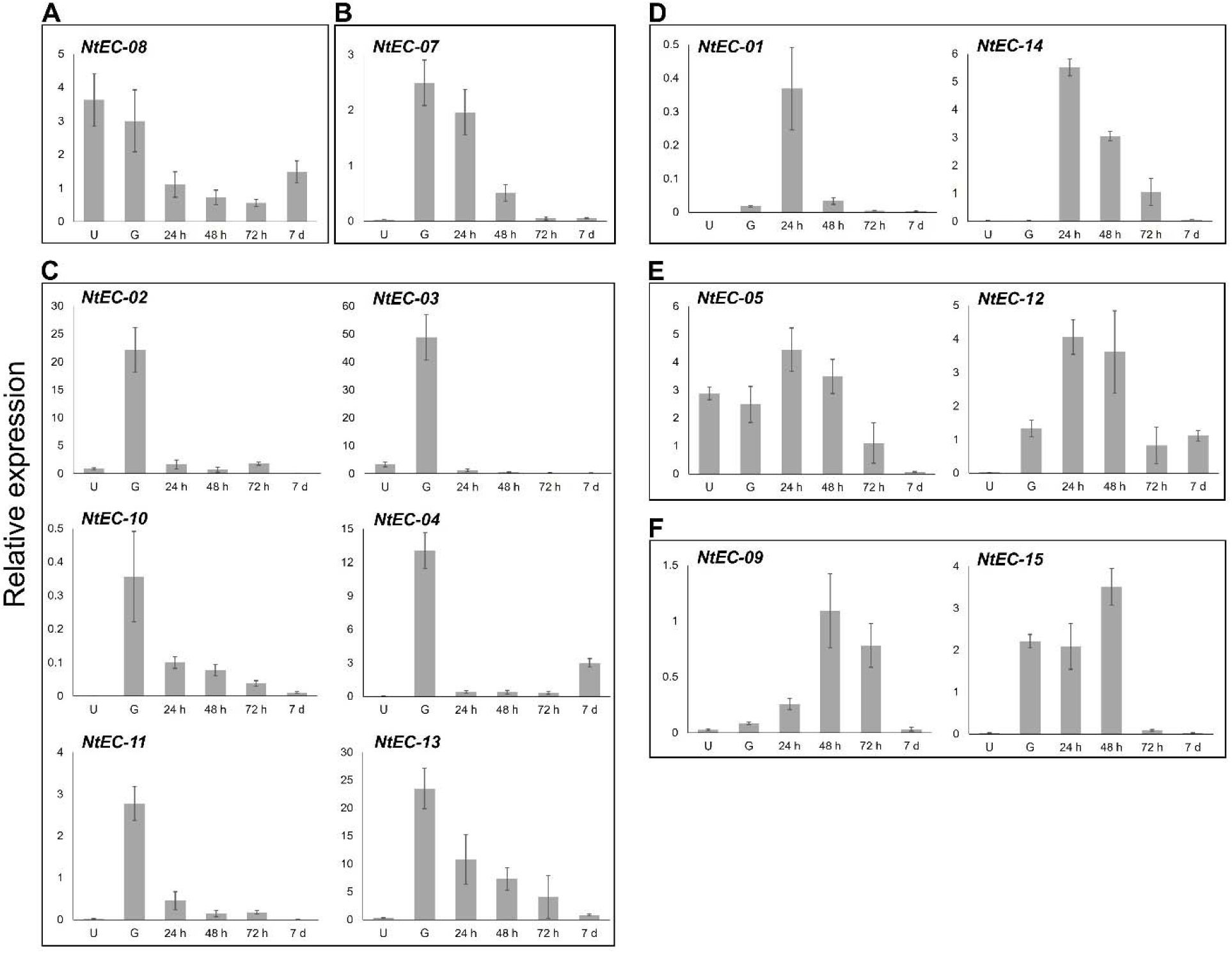
Expression profiles of selected *Neophysopella tropicalis* effector candidate genes (*NtECs*) identified from the germinated urediniospore transcriptome. Transcript accumulation of *NtECs* was assessed by real-time quantitative PCR (RT-qPCR) in samples obtained from dormant urediniospores (U), germinated urediniospores on polystyrene plates (G), and infected *Vitis labrusca* cv. Nagara Rosada leaves at 24, 48, and 72 hours post-inoculation (hpi), and 7 days post-inoculation (dpi). Genes were grouped into panels (A–F) based on shared expression patterns: (A) *NtEC* with highest expression in dormant and germinated urediniospores; (B) *NtEC* with highest peaks in germinated urediniospores and at 24 hpi; (C) *NtECs* with peak expression exclusively in germinated urediniospores; (D) *NtECs* with highest expression at 24 hpi; (E) *NtECs* peaking at 24 and 48 hpi; (F) *NtECs* with peak expression at 48 hpi. Bars represent mean relative expression levels, normalized using the mean transcript accumulation of four *N. tropicalis* endogenous genes: β-tubulin 1 (*Nt*β*Tub1*), β-tubulin 2 (*Nt*β*Tub2*), glyceraldehyde-3-phosphate dehydrogenase (*NtGAPDH*), and cytochrome c oxidase subunit III (*NtCytIII*). Error bars indicate the standard error of the mean (SEM). The primer sequences are listed in Table S3.

### Microscopy analyses of the compatible *Vitis labrusca*–*Neophysopella tropicalis* interaction

To visualize the grapevine rust fungus, infected host leaf samples were stained with wheat germ agglutinin (WGA) conjugated to fluorescein isothiocyanate (FITC), a lectin that binds specifically to chitin, a major component of the fungal cell wall (Marrafon-Silva et al., 2024). Microscopy analysis showed that *N. tropicalis* had reached the penetration stage at 24 hpi, as evidenced by the formation of elongated germ tube terminating in well-defined appressorium (Fig. 4A). At this time point, fungal hyphae were also observed penetrating host tissue through stomatal openings (Fig. 4B). Most germ tubes had developed appressoria at 24 hpi (Fig. 4C–E), and similar infection structures remained evident at 48 hpi (Fig. 4F, G). Notably, intracellular infection hyphae were visible within host tissues at 48 hpi (Fig. 4H, I). Based on our experimental conditions, these findings suggest that *N*. *tropicalis* completes the penetration phase and transitions into the early stage of host colonization by 48 hpi.

**Fig. 4:**
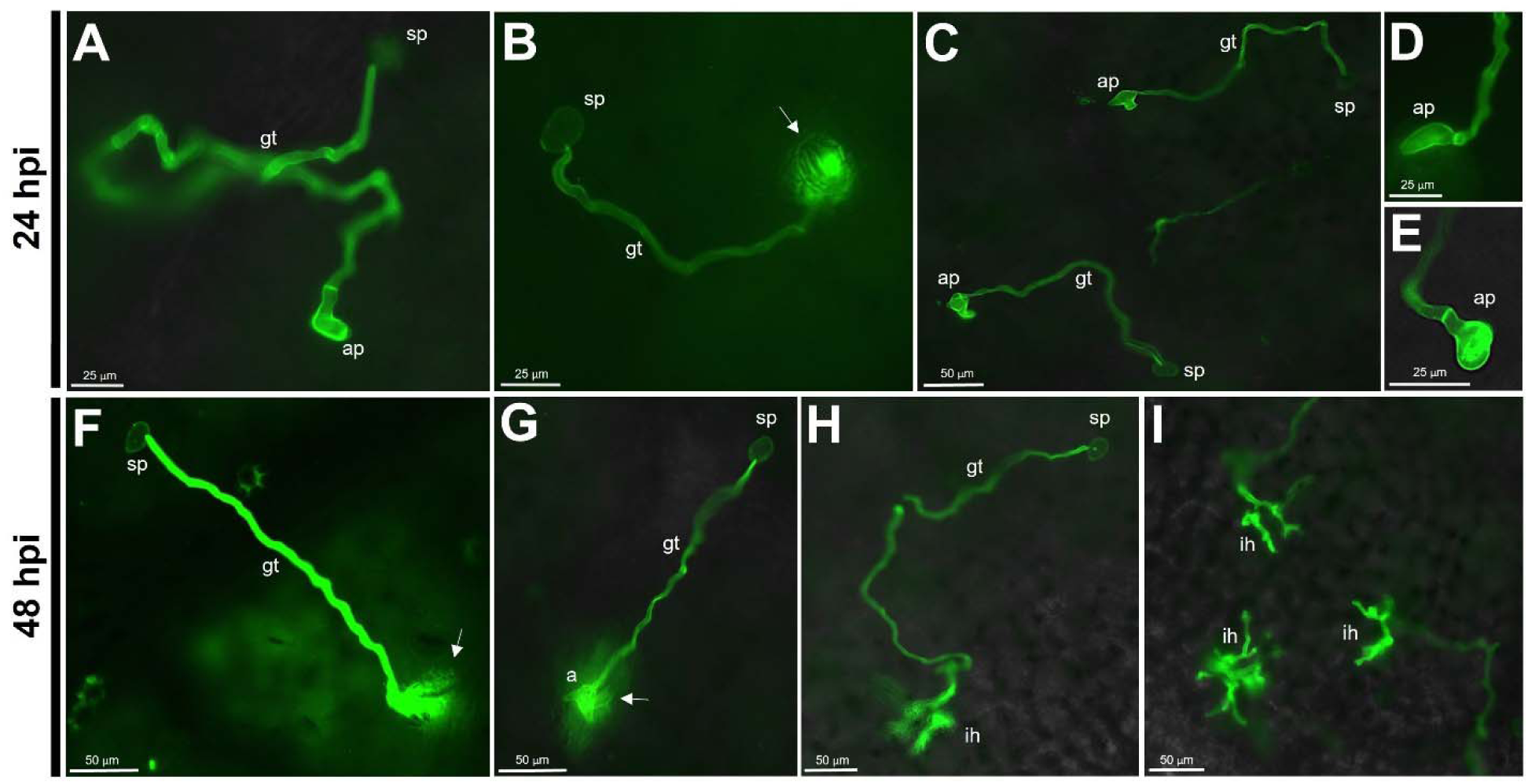
*Neophysopella tropicalis* development during early phases of infection. (A-E) Penetration phase at 24 hours post-inoculation (hpi). (F-I) Transition from penetration to early colonization at 48 hpi. Fully expanded leaves of *Vitis labrusca* cv. Niagara Rosada were inoculated with 1 x 10^5^ uredinipospores/mL. After 24 hours of incubation in a dark, humid chamber, the plants were transferred to a controlled environment with a 12 h light/12 h dark photoperiod at 25 °C. Fungal structures: urediniospores (sp), germ tubes (gt), appressoria (ap), and infection hyphae (ih) are stained green with WGA-FITC. Time points are indicated by the black lines on the left. Stomata are indicated by white arrows.

At 72 hpi, infection hyphae began proliferating within the grapevine leaf tissue, marking the onset of a well-established colonization stage (Fig. 5A–C). By 5 dpi, extensive radial growth of intercellular hyphae was observed, forming broad hyphal networks at the infection sites (Fig. 5D). At this point, the coalescence of hyphal networks originating from individual urediniospores is also evident (Fig. 5E). Based on these observations, the colonization phase of *N*. *tropicallis* was noticed as occurring between 72 hpi and 5 dpi.

**Fig. 5:**
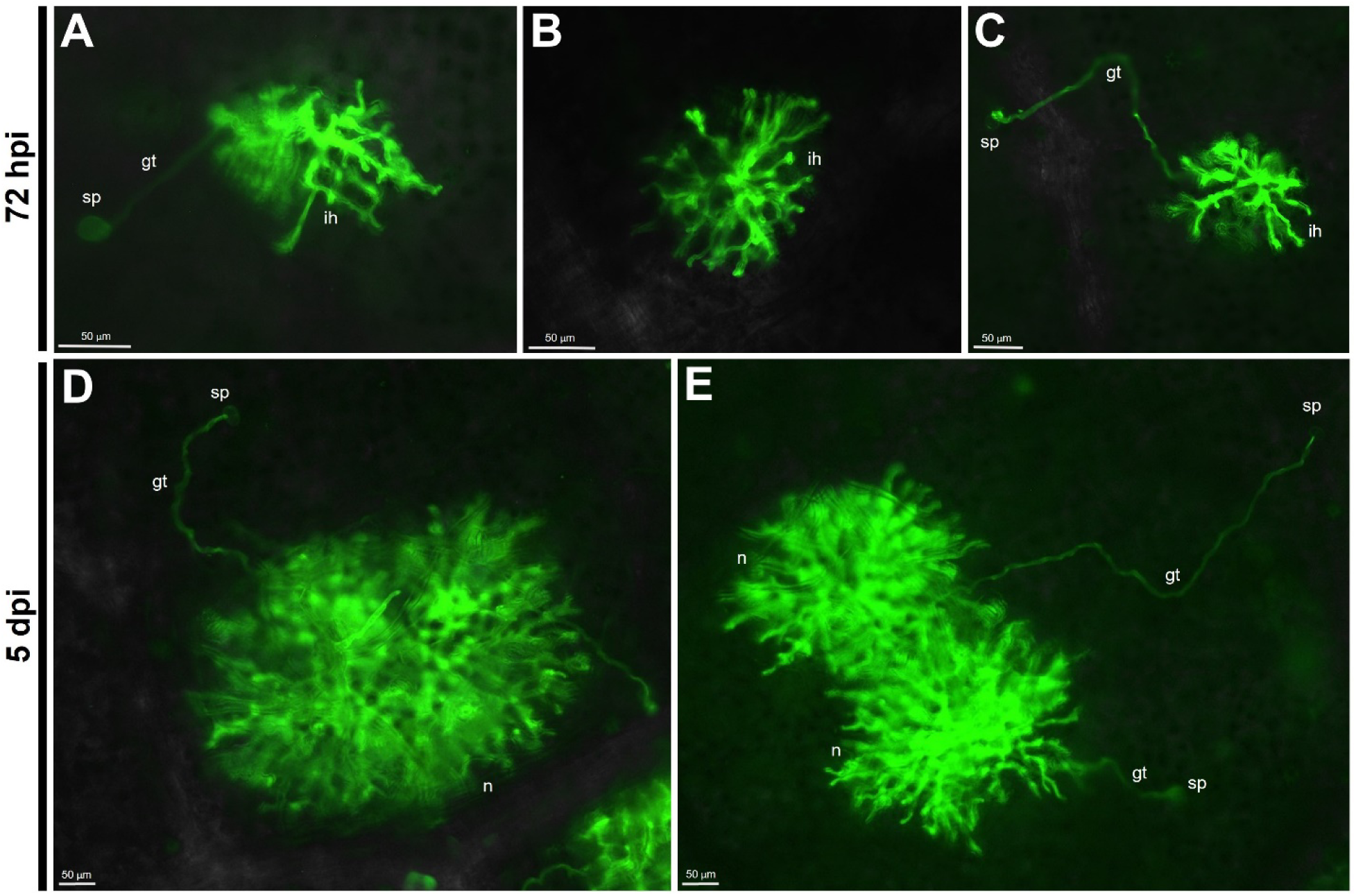
*Neophysopella tropicalis* colonization phase of infection. (A-C) Infected leaves at 72 hours post-inoculation (hpi). (D-E) Infected leaves at 5 days post-inoculation (dpi). Fully expanded leaves of *Vitis labrusca* cv. Niagara Rosada were inoculated with a suspension containing 1 × 10 urediniospores/mL. Following 24 hours of incubation in a dark, humid chamber, the plants were transferred to a controlled environment maintained at 25 °C with a 12-hour light/12-hour dark photoperiod. Fungal structures, including urediniospores (sp), germ tubes (gt), appressoria (ap), infection hyphae (ih), and hyphal network (n), were visualized by staining with WGA-FITC, appearing green under fluorescence. A black lines on the left denote each time point.

By 7 hpi, hyphal networks had further expanded, and both closed and partially opened uredia containing urediniospores were observed (Fig. 6A). At 9 dpi, all rust pustules exhibited fully mature, open uredia abundant in urediniospores (Fig. 6B, C), marking the reproductive phase of *N. tropicalis*. Having established a clear timeline for each infection stage, from penetration (24– 48 hpi), through colonization (72 hpi to 5 dpi), and up to reproduction (7–9 dpi), we can now correlate these developmental phases with the transcript accumulation profiles of individual *NtEC* genes. The next step is to functionally characterize these candidates to identify those that act as bona fide effectors.

**Fig. 6:**
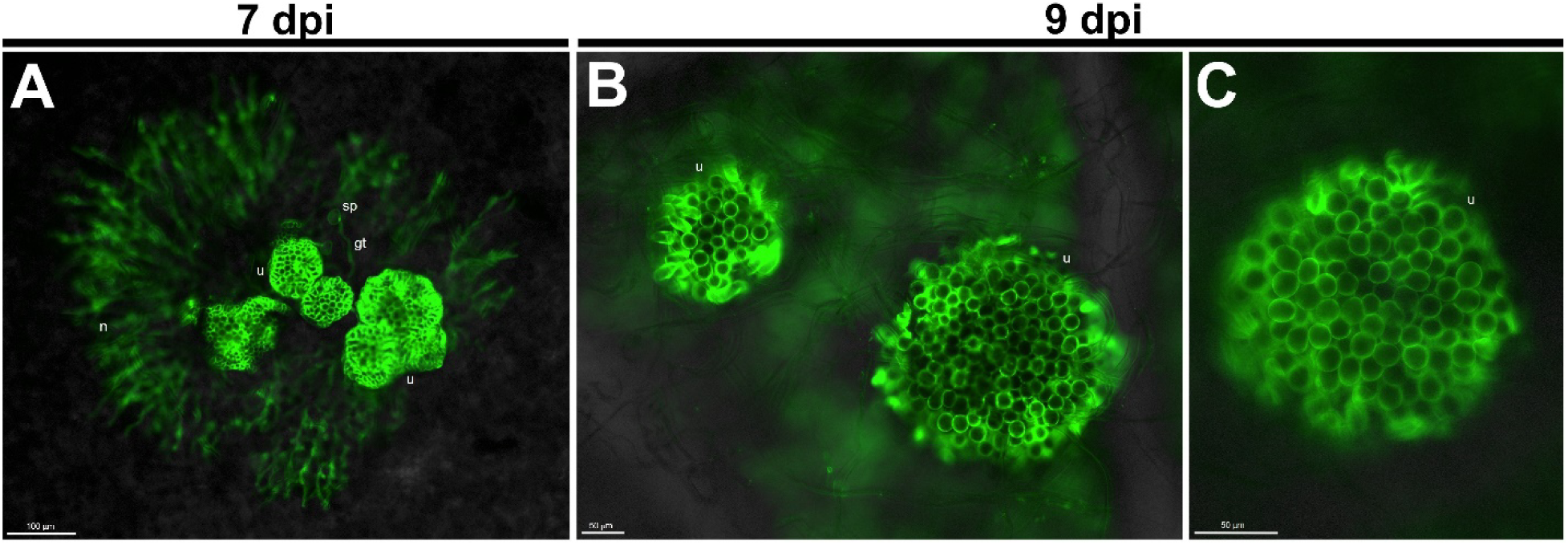
*Neophysopella tropicalis* reproductive phase. (A) Infected leaves at 7 days post-inoculation (dpi). (B, C) Infected leaves at 9 dpi. Fully expanded leaves of *Vitis labrusca* cv. Niagara Rosada were inoculated with a suspension containing 1 × 10 urediniospores/mL. Following 24 hours of incubation in a dark, humid chamber, the plants were transferred to a controlled environment maintained at 25 °C with a 12-hour light/12-hour dark photoperiod. Fungal structures—including urediniospores (sp), germ tubes (gt), hyphal network (n), and uredinia (u)—were visualized by staining with WGA-FITC (green fluorescence). Time points are indicated by the black lines at the top.

### *Neophysopella tropicalis* effectors with plant immunity-suppressing activity

To overcome the challenges of functional characterization in obligate biotrophic fungi such as rust pathogens, we employed a Type III Secretion System (T3SS)-mediated delivery method using *Pseudomonas fluorescens* EtHAn. This approach enabled us to individually test the ability of NtECs to suppress plant immune response. Specifically, we tested their capacity to inhibit AvrB-triggered cell death in *Nicotiana benthamiana* leaves (Maia et al., 2022).

Nine *NtEC* coding sequences (NtEC-03, -05, -09, -10, -11, -12, -13, -14 and -15), excluding their predicted native signal peptides, were cloned into the pEDV6 vector, and each pEDV6::*NtEC* construct was mobilized into *Pseudomonas fluorescens* EtHAn for in planta delivery. Six of the nine candidates tested were able to suppress AvrB-induced cell death (Fig. 7A). Among these, three effectors (NtEC-05, NtEC-09, and NtEC-10) exhibited moderately high suppression activity, with consistent suppression frequencies ranging from 53% to 66% across three independent experiments (Fig. 7B; Table S4). In addition, three effectors (NtEC-11, NtEC-12, and NtEC-13) showed moderate suppression activity, ranging from 40% to 46% (Fig. 7B; Table S4). The remaining candidates (NtEC-03, NtEC-14, and NtEC-15) displayed suppression levels below 30% and were therefore considered to lack cell death suppression activity in our assay (Supplementary Fig. S5).

**Fig. 7:**
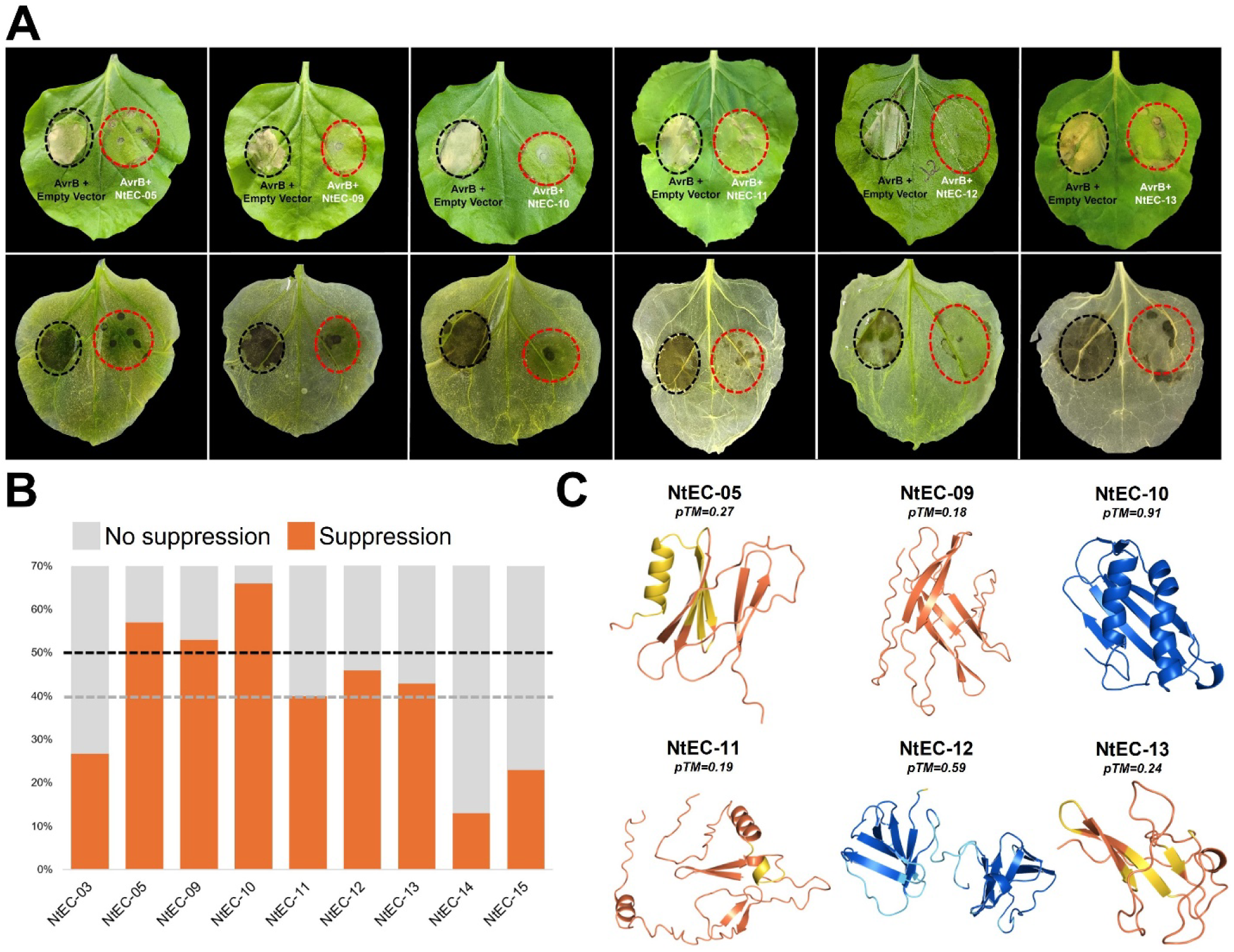
*Neophysopella tropicalis* effectors exhibiting cell death suppression activity. (A) Six effectors suppress tissue collapse associated with AvrB-induced cell death in *Nicotiana benthamiana*. Bacterial suspensions of *Pseudomonas fluorescens* EtHAn co-expressing pVSP::*AvrB* and the empty vector pEDV6 (AvrB + empty vector) were mixed in a 1:1 ratio to a final concentration of 2 × 10 CFU mL ¹ and used as a negative control (black dotted circle). For experimental treatments, *P. fluorescens* EtHAn strains expressing pVSP::*AvrB* were co-infiltrated with strains individually expressing each candidate effector (AvrB + NtEC) under the same conditions (red dotted circle). The upper panel shows leaf images captured 72 hours post-infiltration, while the lower panel displays the same leaves after ethanol bleaching to enhance visualization of cell death. (B) Histogram showing the extent of cell death suppression observed for nine different pEDV6::*NtEC* constructs. Suppression levels were determined based on the percentage frequency across all co-infiltrated leaves, derived from three independent experiments with ten biological replicates each (see Supplementary Table S4). Suppression levels greater than 50% (black dashed line) were categorized as moderately high, whereas levels above 40% (gray dashed line) were regarded as moderate. (C) Structural predictions of the six NtEC cell death suppressors using AlphaFold v3. Structures are color-coded based on pLDDT scores: high confidence (dark blue), medium confidence (cyan), low confidence (orange), and very low confidence (yellow). The predicted TM-score (*pTM*) is shown for each model.

Furthermore, the three-dimensional structures of the six cell death-suppressing NtECs were predicted using AlphaFold v3 (Abramson et al., 2024). Low-confidence predictions (pTM < 0.5) were obtained for NtEC-05, NtEC-09, NtEC-11, and NtEC-13 (Fig. 7C), while NtEC-10 and NtEC-12 yielded high-confidence models (pTM > 0.5). Notably, NtEC-10 exhibited both high structural similarity and significant sequence identity with a predicted glycan-binding protein Y3-like domain-containing protein from *Phakopsora pachyrhizi* (UniProt ID: A0A0S1MKU5) (Supplementary Fig. S6). Together, these results highlight a subset of *N. tropicalis* effectors with potential roles in host immunity suppression and offer initial structural insights to guide future functional studies.

## Discussion

To our knowledge, this is the first transcriptome analysis of *N. tropicalis*. We focused on the urediniospore germination stage because it represents a key moment when the fungus initiates the infection process (Shen et al., 2017; Lorrain et al., 2019). By examining this stage, we identified candidate effector proteins that are likely important for the establishment of infection and colonization of host tissue. From more than 64,000 assembled transcripts, we predicted 386 secreted proteins with features typical of fungal effectors. This dataset considerably expands our understanding of the *N. tropicalis* secretome and provides a collection of high-confidence effector candidates suitable for functional assays to identify bona fide effectors with immune-suppressing activity and broader relevance to grapevine–rust interactions.

In addition to identifying secreted protein genes, our transcriptome assembly revealed full-length ORFs for a second β-tubulin gene (*Nt*β*Tub2*), not previously reported in *N. tropicalis*, as well as for a glyceraldehyde-3-phosphate dehydrogenase (*NtGAPDH*) gene. The availability of the complete β-tubulin sequence is particularly relevant because this gene is a well-known target of benzimidazole fungicides, and access to its full coding region will allow monitoring of mutations associated with fungicide resistance (Zhao et al., 2014). Beyond this applied relevance, both β*-tubulin* and *GAPDH* also serve as reliable endogenous reference genes, making them useful for accurate normalization in RT-qPCR expression studies.

From the 386 *N. tropicalis* candidate effectors (NtECs), we selected fifteen highly expressed in germlings and experimentally validated fourteen by RT-PCR. These were further characterized by their expression patterns at different stages of *N. tropicalis* pathogenesis using RT-qPCR. The results revealed diverse temporal expression profiles, suggesting functional differentiation within the effector repertoire (Jaswal et al., 2021). Eight *NtECs* (e.g., *NtEC-02*, -*03*, -*04*, -*07*, -*08*, -*10*, - *11*, -*13*) were preferentially expressed in germinated urediniospores, pointing to roles in pre-biotrophic development and preparation for host entry. Others, such as *NtEC-01* and *NtEC-14*, showed peak expression exclusively at 24 hpi, consistent with a function in penetration and appressorium-associated processes. *NtEC-05* and *NtEC-12* displayed sustained expression across early stages, suggesting functions that extend from penetration through the initial establishment of biotrophy. Finally, *NtEC-09* and *NtEC-15* reached maximal expression at 48 hpi, indicating a role in sustaining the early stage of colonization. To further support these temporal expression patterns, we coupled gene expression profiling with WGA-FITC staining microscopy (Marrafon-Silva et al., 2024).

At 24 hpi, *N. tropicalis* had developed fully differentiated appressoria with clear septation and initiated host penetration, corresponding tightly with the peak expression of five *NtEC* genes (*NtEC-01*, -*05*, -*07*, -*12*, -*14*). Microscopy revealed that appressoria frequently formed directly above stomatal openings, suggesting that *N*. *tropicalis* may exploit these natural openings in grapevine leaves as entry points into host tissues. Stomata, however, are not merely passive entry sites but function as dynamic immune structures capable of perceiving microbial signals and activating defense responses (Melotto et al., 2024). Guard cells recognize fungal cell-wall-derived chitin via the CERK1 receptor complex, triggering calcium influx, activation of S-type anion channels, and rapid stomatal closure to restrict pathogen ingress (Ye et al., 2020). Some fungi, however, circumvent stomatal immunity by converting chitin to chitosan oligosaccharides or secreting effectors that interfere with guard-cell signaling, thereby preventing closure or inducing guard-cell death. The high expression of these five *NtECs* during penetration suggests that they may contribute to the suppression of guard-cell–mediated defenses or alteration of stomatal signaling pathways, facilitating *N*. *tropicalis* entry through these openings. By interfering with stomatal immunity at this early stage, *N*. *tropicalis* likely achieves efficient penetration while minimizing host recognition and the energetic cost of direct cuticle degradation. Following successful entry, infection progressed rapidly: by 48 hpi, small infection hyphae were visible, marking the transition from penetration to early colonization and coinciding with maximal expression of *NtEC-09* and *NtEC-15*. These findings demonstrate that germinated urediniospores produced *in vitro* represent a valuable biological source for identifying *N*. *tropicalis* candidate effector genes expressed during the initial stages of infection.

Although no candidate effector was preferentially induced at later stages, WGA-FITC staining revealed that between 72 hpi and 5 dai, the fungus entered an active colonization phase characterized by extensive proliferation of infection hyphae within host tissues. By 7–9 dai, fungal growth was widespread, with expanding hyphal networks giving rise to rust pustules containing uredia. Together, these histological observations offer an unprecedented timeline reported so far of the *N. tropicalis–V. labrusca* compatible interaction and establish a clear framework for interpreting candidate effector gene expression profiles in the context of fungal development.

A significant advance of this study was the functional analysis of NtEC proteins, where we tested their ability to suppress AvrB-induced cell death in *N. benthamiana* leaves using *P. fluorescens* EtHAn-adapted effector detector vector system (Maia et al., 2022). Six effectors (NtEC-05, -09, - 10, -11, -12, -13) successfully suppressed AvrB-triggered cell death, with NtEC-05, NtEC-09, and NtEC-10 displaying the strongest activity. NtEC-05 appears to be specific to the order Pucciniales, NtEC-10 is broadly conserved across Basidiomycota and Ascomycota, and NtEC-09 showed no homologs, suggesting that it may be unique to *N. tropicalis*. The capacity of numerous NtECs to suppress effector-triggered immunity supports the idea that *N. tropicalis* deploys a redundant and multilayered effector repertoire to ensure successful infection. These findings parallel functional studies in *Phakopsora pachyrhizi*, *Hemileia vastatrix*, and *Sporisorium scitamineum*, where bacterial Type Three Secretion System-based delivery systems have uncovered effectors that modulate cell death responses, underscoring the value of surrogate approaches for investigating obligate biotrophs (Qi et al., 2016; Maia et al., 2017; Teixeira-Silva et al., 2020; Maia et al., 2022).

Structural predictions generated by AlphaFold v3 provide preliminary clues into *N. tropicalis* effector architecture. Although some models (NtEC-05, -09, -11, -13) yielded low-confidence outputs, NtEC-10 and NtEC-12 produced robust 3D predictions. Notably, NtEC-10 exhibited strong similarity to a glycan-binding protein domain predicted in *P. pachyrhizi*, suggesting a conserved biochemical role possibly linked to host cell wall interactions (Tanaka and Kahmann, 2021). These predicted folds also offer a basis for comparative analyses with effectors from other rust fungi, enabling the identification of conserved structural motifs or domains associated with immune suppression (Le Naour-Vernet et al., 2025).

Overall, this work provides a first step toward integrating effector biology into breeding for rust resistance in grapevine, extending strategies that have proven effective against other viticultural diseases (Paineau et al., 2025). The *N*. *tropicalis* effectors identified here could serve as molecular probes to screen grapevine germplasm for recognition-based resistance, accelerating the development of effector-assisted breeding programs. Cross-species comparisons of effector repertoires may further reveal conserved virulence determinants that could act as durable resistance targets. As for future studies, the directions could be: (i) identifying the host targets of NtECs to elucidate their molecular mechanisms, (ii) validating their functions directly in grapevine through transient expression assays, and (iii) expanding genomic resources for *N. tropicalis* to contextualize effector evolution better. Collectively, these efforts will deepen knowledge of grapevine–rust interactions and advance the development of sustainable control strategies.

## Author contributions

TM conceived and designed the study. TM and JVFM methodology. RGHB and TM data analysis. TM wrote and edited the manuscript with input from CBM-V. CBM-V funding acquisition.

## Data availability

The datasets generated and analyzed during the current study are available in the NCBI BioProject database (http://www.ncbi.nlm.nih.gov/bioproject/) under accession PRJNA1346437. The associated BioSamples are SAMN52835188, SAMN52835189, and SAMN52835190.

## Acknowledgments

We thank Elaine Vidotto Batista (Genomics Group Lab, ESALQ/USP) for her excellent lab. technical assistance, and the group of Professor Lilian Amorim (Department of Plant Pathology, ESALQ-USP) for their support with the grapevine rust fungus and grapevine plants.

## Supplementary Figures & Legends

**Fig. S1:**
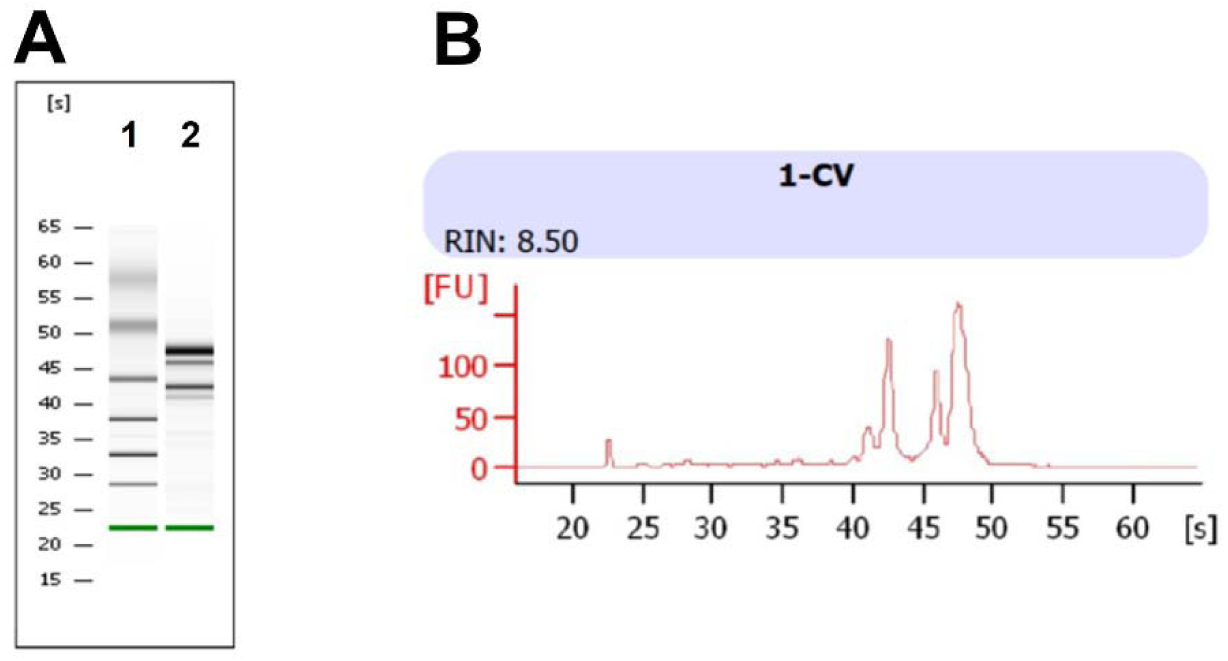
Evaluation of RNA quality from germinated urediniospores of *Neophysopella tropicalis*. (A) Capillary electrophoresis analysis enabled visual evaluation of RNA integrity. Lane 1: RNA ladder; Lane 2: total RNA extracted from germinated urediniospores. (B) The corresponding electropherogram displays well-defined 18S and 28S rRNA peaks, indicative of high-quality RNA. The RNA Integrity Number (RIN) exceeded 8.0, confirming that the sample is of excellent quality for downstream high-throughput sequencing.

**Fig. S2.**
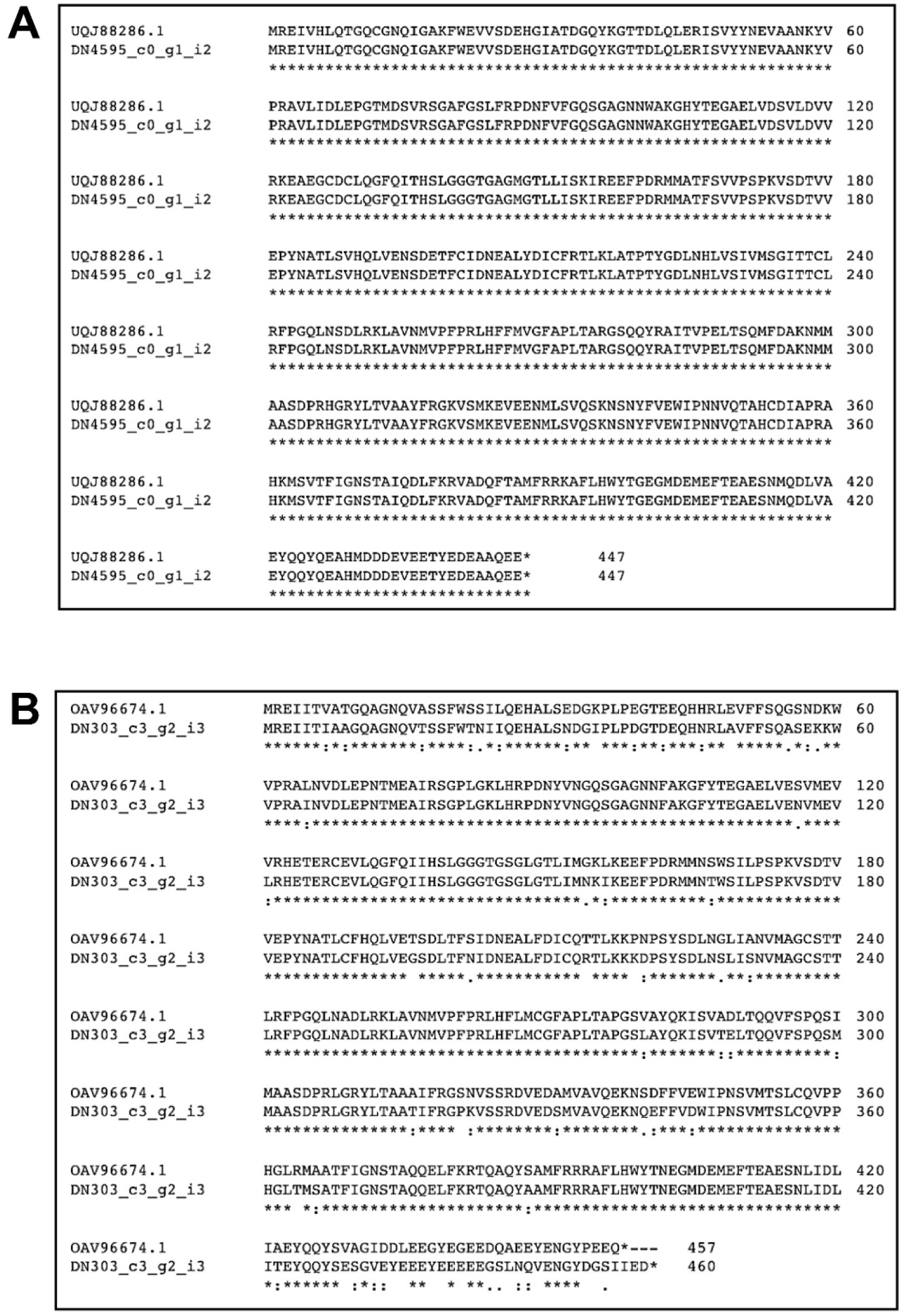
Alignment of deduced amino acid sequences of two β-tubulin genes identified in the *Neophysopella tropicalis* transcriptome. **(A)** Alignment of the full-length ORF represented by contig DN4595_c0_g1_i2 (*Nt*β*Tub1*) with the *N. tropicalis* β-tubulin protein (GenBank accession: UQJ88286.1), the top BLASTP hit (E-value = 0.0). **(B)** Alignment of the full-length ORF represented by contig DN303_c3_g2_i3 (*Nt*β*Tub2*) with the *Puccinia triticina* β-tubulin protein (GenBank accession: OAV96674.1), the top BLASTP hit (E-value = 0.0). Alignments were generated using Clustal Omega. Markers: “*” = identical residues; “:” = conserved substitutions; “.” = semi-conserved substitutions.

**Fig. S3.**
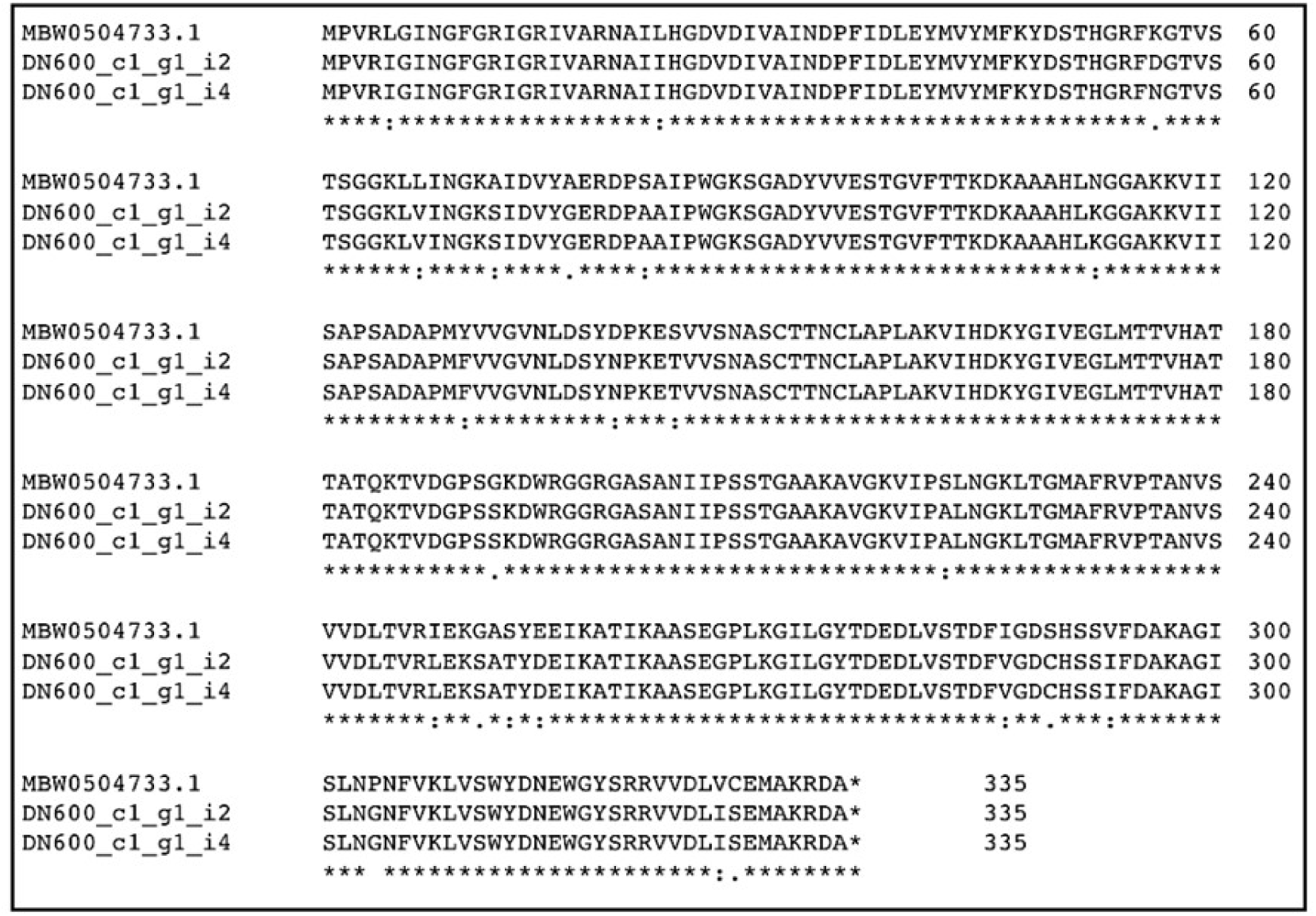
Alignment of the deduced amino acid sequences from two full-length ORFs, represented by contigs DN600_c1_g1_i2 and DN600_c1_g1_i4, with the glyceraldehyde-3-phosphate dehydrogenase (GAPDH) protein from *Austropuccinia psidii* (top BLASTP hit; accession: MBW0504733.1; E-value = 0.0). The alignment was performed using Clustal Omega. Symbols indicate: “*” = identical residues; “:” = conserved substitutions; “.” = semi-conserved substitutions.

**Fig. S4.**
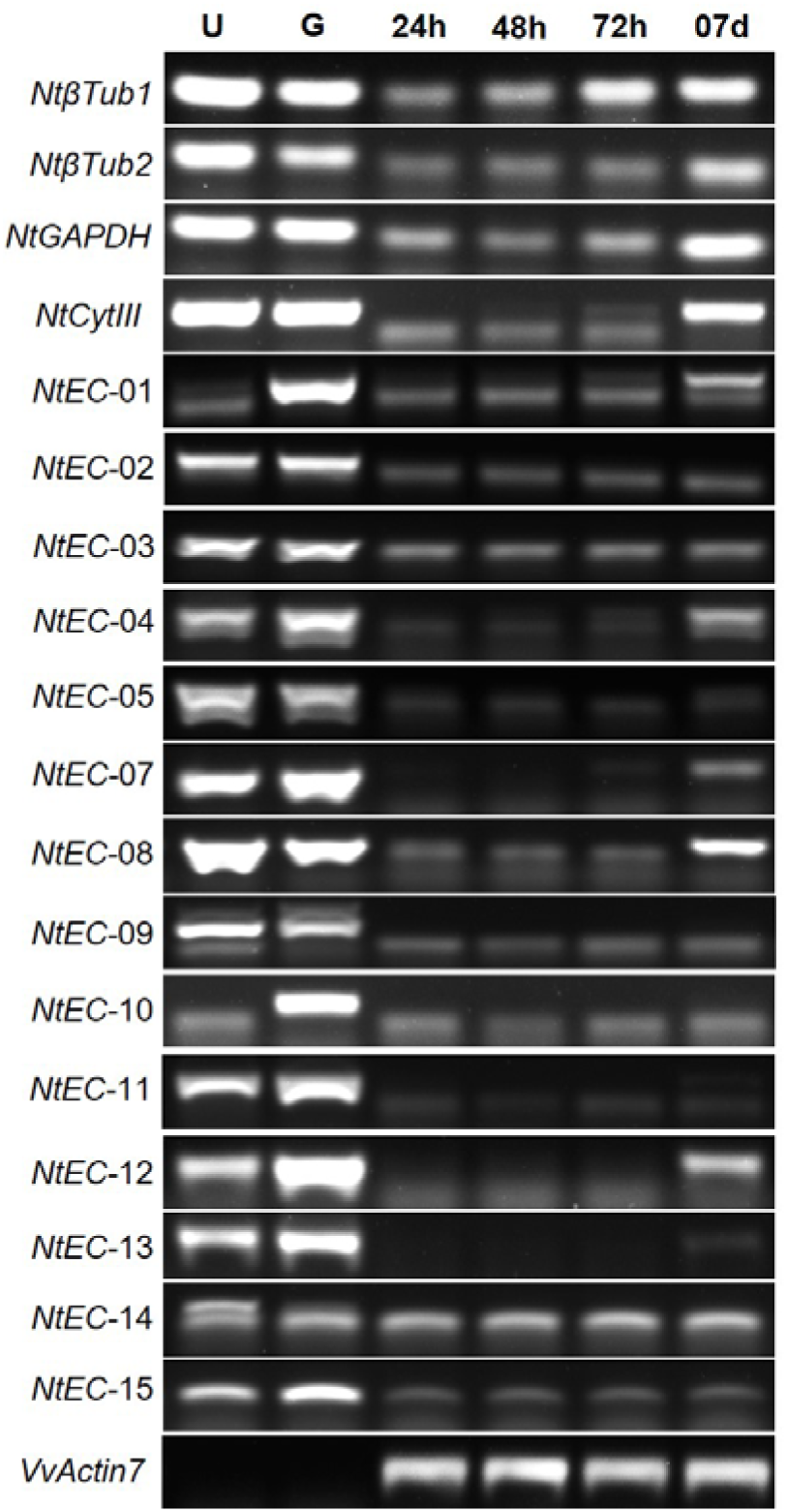
Validation of cDNA synthesis. Reverse transcription-polymerase chain reaction (RT-PCR) amplification of *NtEC* genes was performed using cDNA synthesized from dormant (U) and germinated (G) urediniospores, as well as from infected *Vitis labrusca* cv. Niagara Rosada leaves collected at 24, 48, and 72 hours, and 7 days post-inoculation with *Neophysopella tropicalis* isolate Aglr64. The *N*. *tropicallis* endogenous genes β*-tubulin* 1 and 2 (*Nt*β*Tub1* and *Nt*β*Tub2*), *glyceraldehyde-3-phosphate dehydrogenase* (*NtGAPDH*), and *cytochrome c oxidase subunit III* (*NtCytIII*) served as positive controls for fungal gene expression, while the actin (*VvActin7*) gene was used as a positive control for grapevine. All real-time qPCR primer sets yielded a single, specific amplification product. In a few reactions, a faint secondary band appeared, representing residual by products from the PCR mixture.

**Fig. S5:**
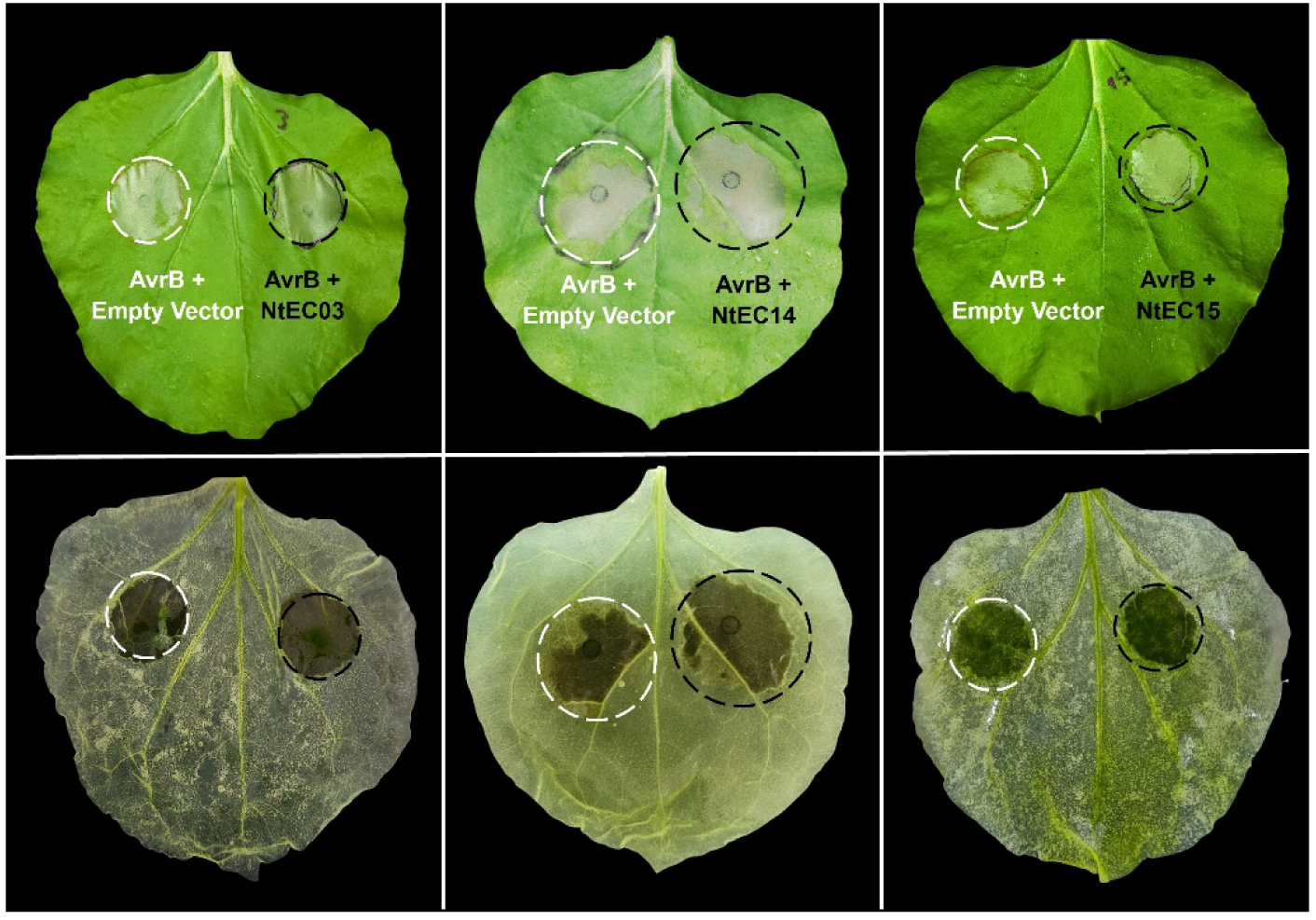
NtEC-03, NtEC-14, and NtEC-015 candidate effectors do not suppress AvrB-induced cell death in *Nicotiana benthamiana* leaves. As a negative control (white dotted circle), *Pseudomonas fluorescens* EtHAn strains co-expressing pVSP::*AvrB* and the empty vector pEDV6 were mixed in equal proportions (1:1) to a final concentration of 2 × 10 CFU mL ¹. For the experimental setup, *P*. *fluorescens* EtHAn strains expressing pVSP::*AvrB* were co-infiltrated with individual strains expressing each candidate effector (AvrB + NtEC) under identical conditions (black dotted circle). The top row presents leaf images taken 72 hours after infiltration, while the bottom row shows the same leaves following ethanol-based clearing to better visualize cell death symptoms.

**Fig. S6:**
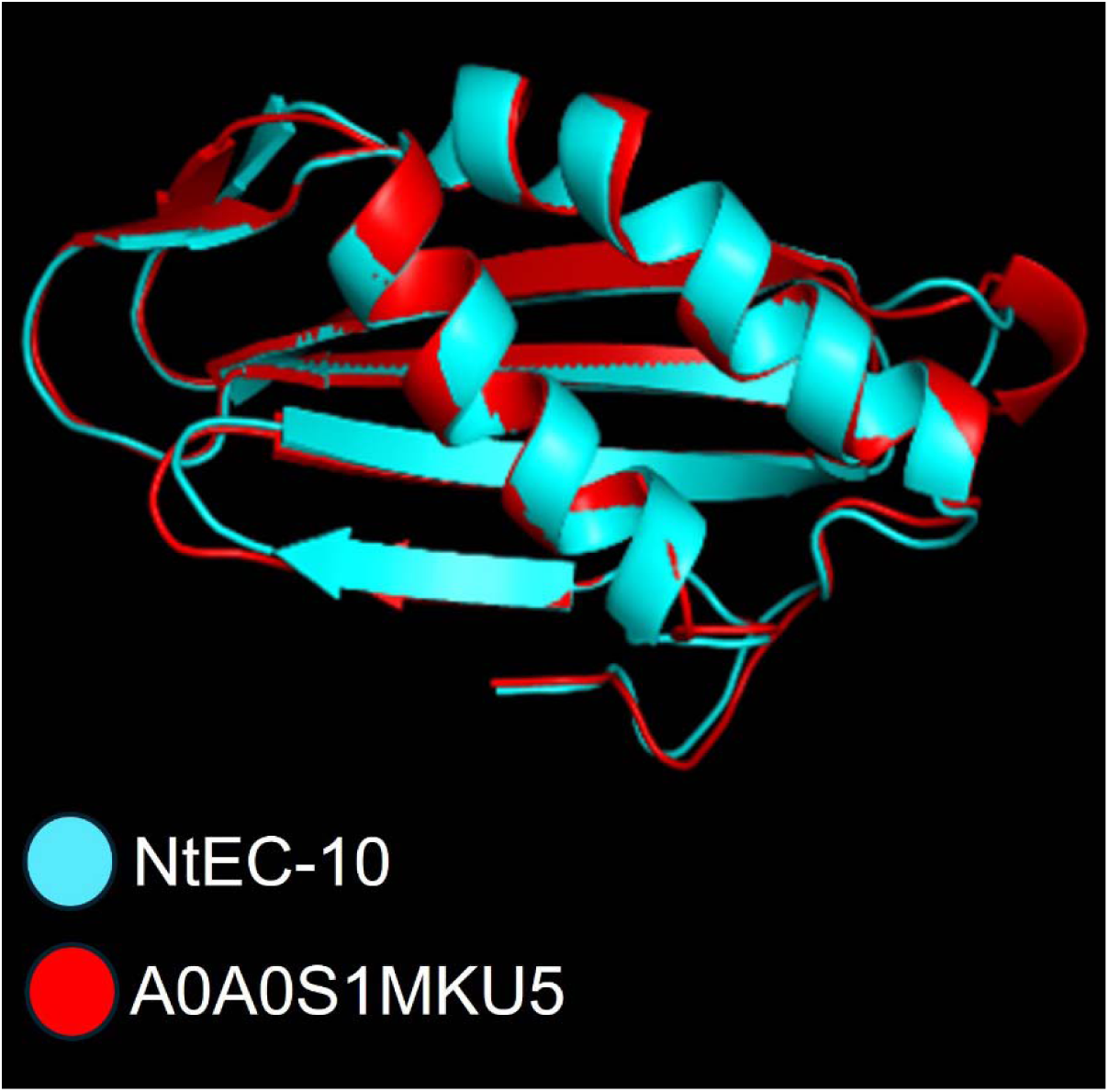
Protein structure alignment of the *Neophysophella tropicalis* effector NtEC-10 with its top structural match. NtEC-10 exhibited strong structural similarity to the predicted glycan-binding protein Y3-like domain-containing protein from *Phakopsora pachyrhizi* (Uniprot ID: A0A0S1MKU5), with TM-score of 0.95520, RMSD of 0.81 Å; and 53% sequence identity. Structural visualization was carried out using PyMOL v3.1.

